# Foundress Number, But Not Queen Size or Boldness, Predicts Colony Life-History in Wild Paper Wasps

**DOI:** 10.1101/609685

**Authors:** Colin M. Wright, David N. Fisher, Wayne V. Nerone, James L.L. Lichtenstein, Elizabeth A. Tibbetts, Jonathan N. Pruitt

**Author notes:** Denotes Corresponding Author.

## Abstract

Colonies of social insects exhibit a spectacular variety of life histories. Here we documented the degree of variation in colony life-history traits, mostly related to productivity, in two species of wild paper wasps. We then tested for associations between colony life-history traits to look for trade-offs or positively associated syndromes, and examined whether individual differences in the behavioral tendencies of foundresses (*Polistes metricus*) or the number of cofoundresses (*P. fuscatus*) influenced colony life-history. The majority of our measures of colony life-history were positively related, indicating no obvious resource allocation trade-offs. Instead, the positive association of traits into a productivity syndrome appears to be driven by differences in queen or microhabitat quality. Syndrome structure differed only marginally between species. Queen boldness and body size were not associated with colony life-history in *P. metricus*. Colonies initiated by multiple *P. fuscatus* foundresses were generally more productive, and this advantage was approximately proportional to the number of cofoundresses. These findings demonstrate that colony life-history traits can be associated together much like individual life-history traits, and the associations seen here convey that differences in overall productivity drive between-colony differences in life-history.

## Introduction

The life-history and behavior of eusocial insects provide some of the most fascinating examples of complex life on Earth (Holldobler & Wilson, 1990). Yet, despite their prominence in evolutionary biology, surprising little is known about the selective forces that have driven the great diversity of colony life-history strategies observed in social insects (Heinze, Kellner & Seal, 2017; Holldobler & Wilson, 1990; Hunt & Toth, 2017; Wcislo & Fewell, 2017). Certain ecological forces, like reduced foundress success due to harsh ecological conditions (e.g., intense competition) (Bartz & Holldobler, 1982; Bernasconi & Strassmann, 1999; Nonacs, 2001) or worker reproduction enabled by longer growth seasons (reviewed in Wcislo & Fewell, 2017) have been rigorously evaluated in the literature. Such studies have enhanced our understanding about the environmental forces that cause group formation in general and some aspects of colony life-history too (e.g., worker reproduction, cofoundress associations). Yet, it is perennially noted that a broader conceptual or theoretical framework to organize the evolution of colony life histories in social insects is still missing (Bourke & Franks, 1995; Heinze et al., 2017; Wcislo & Fewell, 2017).

Intraspecific variation provides evolutionary biologists with a useful tool to probe the selective forces that underlie the evolution of colony life-history traits and to examine how behavior shapes them (Bengston & Jandt, 2014; Jandt, Bengston, Pinter-Wollman, Pruitt, Raine, Dornhaus & Sih, 2014; Wright, Lichtenstein, Doering, Pretorius, Meunier & Pruitt, 2019). By comparing the life-history metrics of natural colonies and testing for associations between them, studies focused on intraspecific variation can provide clues as to which colony life-history traits exhibit the most variability, and whether there are conspicuous associations between them, which could hint at the presence of trade-offs or illuminate possible coadaptation between colony traits (e.g., behavior, demography, nest attributes). Studies conducted on solitary organisms have recovered support for links between life-history trade offs and behavior in some cases (e.g., Niemela, Vainikka, Hedrick & Kortet, 2012; Santostefano, Wilson, Niemela & Dingemanse, 2017). Recent studies on acorn ants (genus *Temnothorax*) have likewise found that variation in resource allocation (i.e., production of workers vs. reproductives) is linked to colony behavior and intraspecific competition in *T. regatulus* (Bengston, 2018; Bengston & Dornhaus, 2015; Bengston, Shin & Dornhaus, 2017). Similar associations appear to present in the related *T. longispinosis* (Segev, Burkert, Feldmeyer & Foitzik, 2017), and there is circumstantial evidence for similar associations in fire ants (Bockoven, Wilder & Eubanks, 2015) and gypsy ants too (Blight, Villalta, Cerda & Boulay, 2016). Yet, whether similar associations are present in other kinds of insect societies, for instance those with weaker reproductive division of labor, is unclear.

In weakly or “primitively” eusocial insect societies (e.g., bumblebees, sweat bees, paper wasps), there are at least two strong candidates for causing intraspecific associations between life-history, behavior, and performance. The first of these are behavioral differences among founding queens in solitary founding species (Wright, Hyland, Izzo, McDermott, Tibbetts & Pruitt, 2018). This is because the initial work and investment of a singleton foundress (e.g., colony establishment, brood care, defence) can strongly influence in a colony’s development, behavior, and functioning (e.g., Wright et al. 2017). This is true of highly eusocial species as well (Fewell & Page, 1999; Shaffer, Sasaki, Haney, Janssen, Pratt & Fewell, 2016), but we predict such effects will diminish as colonies grow and develop in highly eusocial species. Second, in multi-queen founding species, the number of foundresses is likely to influence colony life-history trajectories (Field, Solis, Queller & Strassmann, 1998; Tibbetts & Reeve, 2000). For such societies, the aid of fellow cofoundresses is likely to be influential in shaping colony life-history, because the burdens of founding a colony can be shared among coalition members (Bernasconi & Strassmann, 1999; Bourke & Franks, 1995).

Here we use co-occurring species of weakly eusocial paper wasps to perform an assessment of colony life-history traits in situ. Specifically, we document the degree of variability in a variety of colony life-history traits including the nest initiation date, length of reproductive period, timing of the peak of reproduction, the height of this peak, and total colony production (workers + reproductives) over the course of a colony generation in the wild. We then test for associations between these life-history traits to probe for possible resource allocation trade-offs that might be consistent with alternative life-history strategies, or for positive associations of traits into productivity syndromes. Finally, we examine whether between-group differences in queen behavior (boldness) and queen body size (head width) are associated with colony life-history patterns in the solitary-nesting *Polistes metricus* [mean_foundress_ = 1 07, 95% CI: 1.06-1.08] (Miller, Bluher, Bell, Cini, da Silva, de Souza, Gandia, Jandt, Loope, Prato, Pruitt, Rankin, Rankin, Southon, Uy, Weiner, Wright, Downing, Gadagkar, Lorenzi, Rusina, Sumner, Tibbetts, Toth & Sheehan, 2018), and whether the number of foundresses influences colony life-history patterns in the facultatively cofounding *P. fuscatus* [mean_foundress_ = 2.16, 95% CI: 1.922.40].

## Methods

### Nest discovery and foundress morphology

On the first week of June 2016, approximately two weeks after these *Polistes* wasps began founding nests, we searched for actively nesting *P. metricus* and *P. fuscatus* on buildings located at the Tuttle Campground, in Linesville, Pennsylvania. This campground has been closed since 2010, and the abandoned structures are prime nesting locations for paper wasps. We located 57 active *P. metricus* and 43 *P. fuscatus* nests. All solitary foundresses were individually captured in order to mark and record their head width to the nearest 0.01mm using digital calipers. We then counted the total number of cells present in the nest at this time. Foundresses were then immediately placed back on their nests. Once all wasps had been measured, we waited 24h before conducting field behavioral assays. Many of the foundresses then abandoned their nests; nest abandonment is relatively common in the early portion of the season in *Polistes*. Thus, the dataset reported herein includes 28 *P. metricus* and 17 *P. fuscatus* nests after removing these abandonment events (Abandonment rates: *P. metricus* = 50.8%, *P. fuscatus* = 60.4%).

### Field boldness assessment

All behavioral trials were conducted between 10:00am and 6:00pm, corresponding to active nesting hours. Boldness, or the propensity to engage in risky behavior (Sloan Wilson, Clark, Coleman & Dearstyne, 1994), was assessed using a method similar to that used in Wright et al (Wright et al., 2018; 2016), but modified to a field setting. It has been shown that queens display repeatable between-queen variation in their willingness to flee from their nests when antagonized, which we used to estimate their boldness. Using a 3-foot long (1cm diameter) wooden dowel, we antagonized each queen by prodding them up to 50 times while they resided on their nests, with a 3s pause in between prods, and recorded how many total prods were required before a queen fled from their nests. Queens that flee after just a few prods are considered to elicit a *shyer* phenotype, while queens that stand their ground after repeated prodding elicit a *bolder* phenotype. The reasoning here is that it is more “risky” for a queen to remain on her nest in the face of danger than it is to fly away and return later when the threat has passed. Queens that never abandoned their nests were given a maximum score of 50. This assay was performed on each queen 4 times over the course of a week, and queens were never tested twice in the same day. Boldness was repeatable (r = 0.582, CIs = 0.363-0.796, p < 0.001) and thus we took the average across the four boldness tests to give a single boldness value per queen. In colonies with multiple foundresses (none for *P. metricus*, 37 of the original 43 for *P. fuscatus*) we could not recognize individuals to conduct repeated assays, and so these foundresses had no boldness scores recorded. Furthermore, as there were multiple queens in these nests, we could not assign a head width to a single individual for this species, and so for these nests head widths were not recorded.

### Colony monitoring

Following behavioral assessment, each colony was checked once a week at 6:00am, when queens are inactive from June 12 to November 12. During these weekly checks, we recorded the total number of wasps on each nest. On November 12 no wasps remained on any nests, and each nest was then collected and brought into the lab where we recorded the final cell count (a proxy of overall colony productivity) and the number of caterpillars discovered of the Pyralid moth parasite *Chalcoela iphitalis*, which commonly parasitizes both *P. metricus* and *P. fuscatus* nests (Nelson, 1968; Starr, 1976). We recorded the following five colony-level life-history traits: the week (scored between 1 and 23) of the first production of a worker, the number of weeks between the first and last incidence of reproduction, the maximum number of new individuals produced in a single week, the week at which this occurred, and the total number of cells at the end of the study.

### Statistical analysis

To determine whether these colony life-history traits were associated into a life-history syndrome, we used structural equation models. These assess the fit of specified covariance structures within the data, allowing one to determine which covariance structure, relating to a hypothesis of interest, is the best supported (Brommer, Karell, Ahola & Karstinen, 2014; Dingemanse, Dochtermann & Wright, 2010; Dochtermann & Jenkins, 2007; Santostefano et al., 2017). Here we compared a model where the five life-history traits were unrelated, to one where they were related into a latent variable of a “life-history syndrome”. Traits can be positively related (e.g. among-colony differences in overall productivity) or negatively related (e.g. trade-offs between reproductive strategies). If the life-history syndrome was better supported, we could then examine the coefficients of each life-history trait to determine how the traits contributed to the syndrome. Following this, we fitted “species” as a grouping variable, and compared that to the model where syndrome structure was the same between the species, to determine whether the syndrome structure differed between the species (summarized in Fig 1).

**Figure 1.**
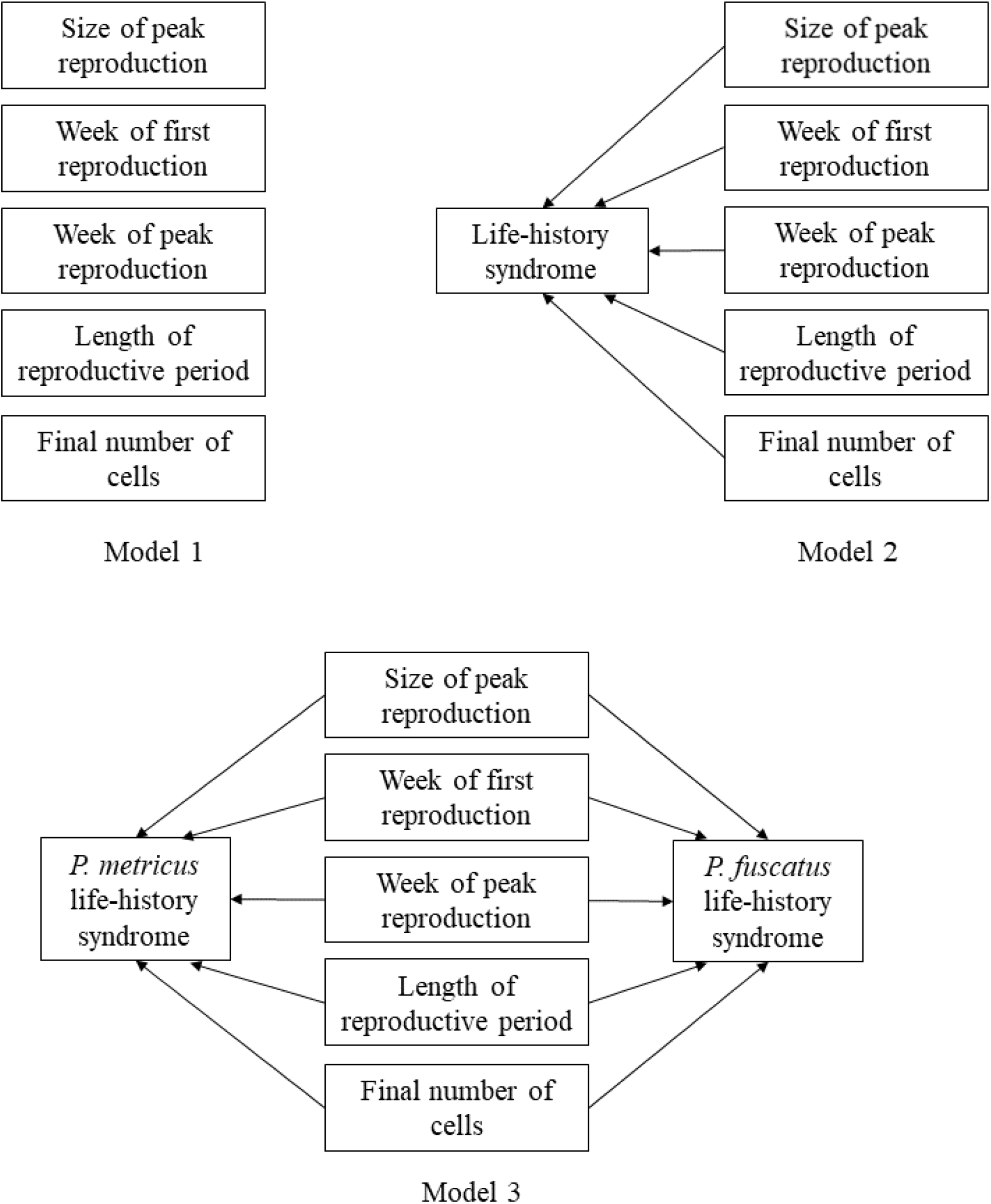
Diagrammatical representations of the three models we compared. In model 1, the five colony life history traits are not related into a syndrome. In model 2, the five traits are related into a life-history syndrome, which is the same across both species. In model 3, the five life-history traits are related in species-specific life-history syndromes.

Structural equation models also allow one to determine whether additional variables influence hypothesized latent variables. For *P. metricus*, we added both foundress head width and average boldness to a model of a *P. metricus*-specific life-history syndrome, to determine whether these traits influence this syndrome. For *P. fuscatus*, we added the number of foundresses to a model of a *P. fuscatus*-specifc life-history syndrome, to determine how this factor influenced the syndrome. We confirmed these results using MANOVAs for each species. Each MANOVA had the five life-history traits as response variables. The MANOVA for *P. metricus* had head width and boldness and predictor variables, while the MANOVA for *P. fuscatus* has number of foundresses as a predictor variable.

## Results

In total, 45 colonies (28 *P. metricus* and 17 *P. fuscatus*) produced at least one worker, and had recordings for: week of first productivity (i.e., worker eclosion), length of reproductive period, timing (week) of maximum productivity, colony growth at maximum productivity, and end number of cells. These were used to test for the presence of a life-history syndrome, and whether it differed between the species. Of these, 18 *P. metricus* had head widths and boldness scores measured, while all *P. fuscatus* nests had the number of foundresses recorded. These factors were used for the species-specific models of what might shape the syndrome.

The model with the life-history syndrome was better supported than the model where the life-history traits were unrelated (model 1 AIC = 1393, model 2 AIC = 1354, ΔAIC = 39). This syndrome differed between the species, with the model with species as a group giving a better fit than the with the same life-history syndrome across species (model 2 AIC = 1354, model 3 AIC = 1327, ΔAIC = 27, Fig 2). The variance-covariance matrix implied by this model is given in Table S1 in the supplementary materials.

**Figure 2.**
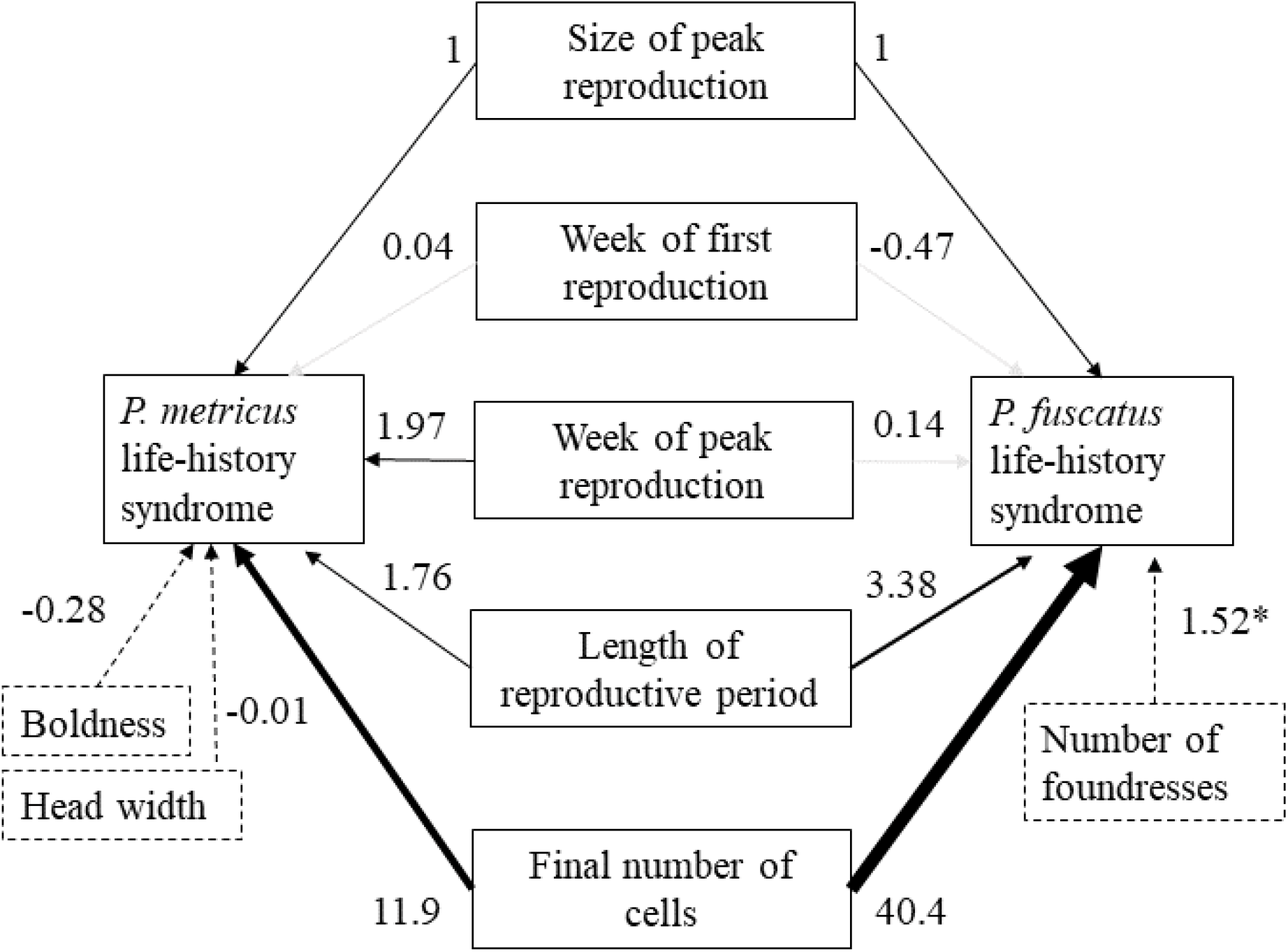
The relative contribution of each life-history trait to the overall life-history syndrome for each species. Numbers are the regression coefficients from the structural equation model. Black lines indicate significant effects, with line thickness representing relative contribution to the life-history syndrome. Grey lines indicate non-significant effects. Note how which traits contribute, and their relative contributions, changes between the species. Also shown are how certain traits (boldness and head width for *P. metricus*, and the number of foundresses for *P. fuscatus*) influence the life-history syndromes. Significant effects here are indicated with an asterix.

In *P. metricus*, longer reproductive periods, later peaks, higher peak productivity, and higher cell counts were all positively associated, while date of first productivity was not related (Table 1, Fig. 2). In *P. fuscatus*, the timing of the productivity peak was not associated into the life-history syndrome, and relative contributions of the remaining three life-history traits to the syndrome differed, but otherwise the overall structure was similar (Table 1, Fig. 2). These suggest a syndrome of overall lower vs. higher productivity, rather than trade-offs between traits.

**Table 1.** Contributions of life-history syndromes in two species of wasp. One trait, here arbitrarily set as maximum productivity, is set as the default, and so the contributions of each trait to the syndrome are relative to this trait’s contribution. Effect estimates are unstandardized.

| Species | Trait | Est (se) | z | p |
| --- | --- | --- | --- | --- |
| *P. metricus* | Maximum productivity | 1 (NA) | NA | NA |
|  | Length of reproductive period | 1.759 (0.812) | 2.168 | 0.030 |
|  | Number of cells at end | 11.923 (3.697) | 3.225 | 0.001 |
|  | Week peak productivity | 1.970 (0.560) | 3.517 | < 0.001 |
|  | First productivity | 0.043 (0.289) | 0.150 | 0.881 |
| *P. fuscatus* | Maximum productivity | 1 (NA) | NA | NA |
|  | Length of reproductive period | 3.383 (1.152) | 2.937 | 0.003 |
|  | Number of cells at end | 40.419 (13.580) | 2.976 | 0.003 |
|  | Week peak productivity | 0.144 (0.780) | 0.185 | 0.853 |
|  | First productivity | -0.474 (0.360) | -1.318 | 0.187 |

In *P. metricus*, neither head width (est = −0.011, SE = 0.524, z = −0.021, p = 0.984) nor average boldness (est = −0.28, SE = 0.021, z = −1.347, p = 0.178) influenced the life-history syndrome (Fig 2). In our MANOVA analysis too, in *P. metricus*, neither head width (MANOVA, Pillai’s Trace_5,11_ = 0.141, approx. F = 0.360, p = 0.865) nor average boldness (MANOVA, Pillai’s Trace_5,11_ = 0.165, approx. F = 0.433, p = 0.816) influenced the life-history syndrome.

In *P. fuscatus*, more foundresses were associated with higher values of this productivity syndrome (est = 1.516, SE = 0.527, z = 2.878, p = 0.004, Figs. 2 & 3). However, there were no differences in per capita measures of life-history traits i.e. after dividing each of the life-history traits by the number of foundresses (est = 0.052, SE = 0.232, z = 0.225, p =0.822). In the parallel MANOVA analysis, more *P. fuscatus* foundresses was associated with higher values of the life-history syndrome (MANOVA, Pillai’s Trace_15_,_33_ = 1.567, approx. F = 2.404, p = 0.018; Fig. 3). However, there were no differences in per capita measures of life-history traits, i.e., after diving each of the life-history traits by the number of foundresses (MANOVA, Pillai’s Trace_15_,_33_ = 1.164, approx. F = 1.394, p = 0.207).

**Figure 3.**
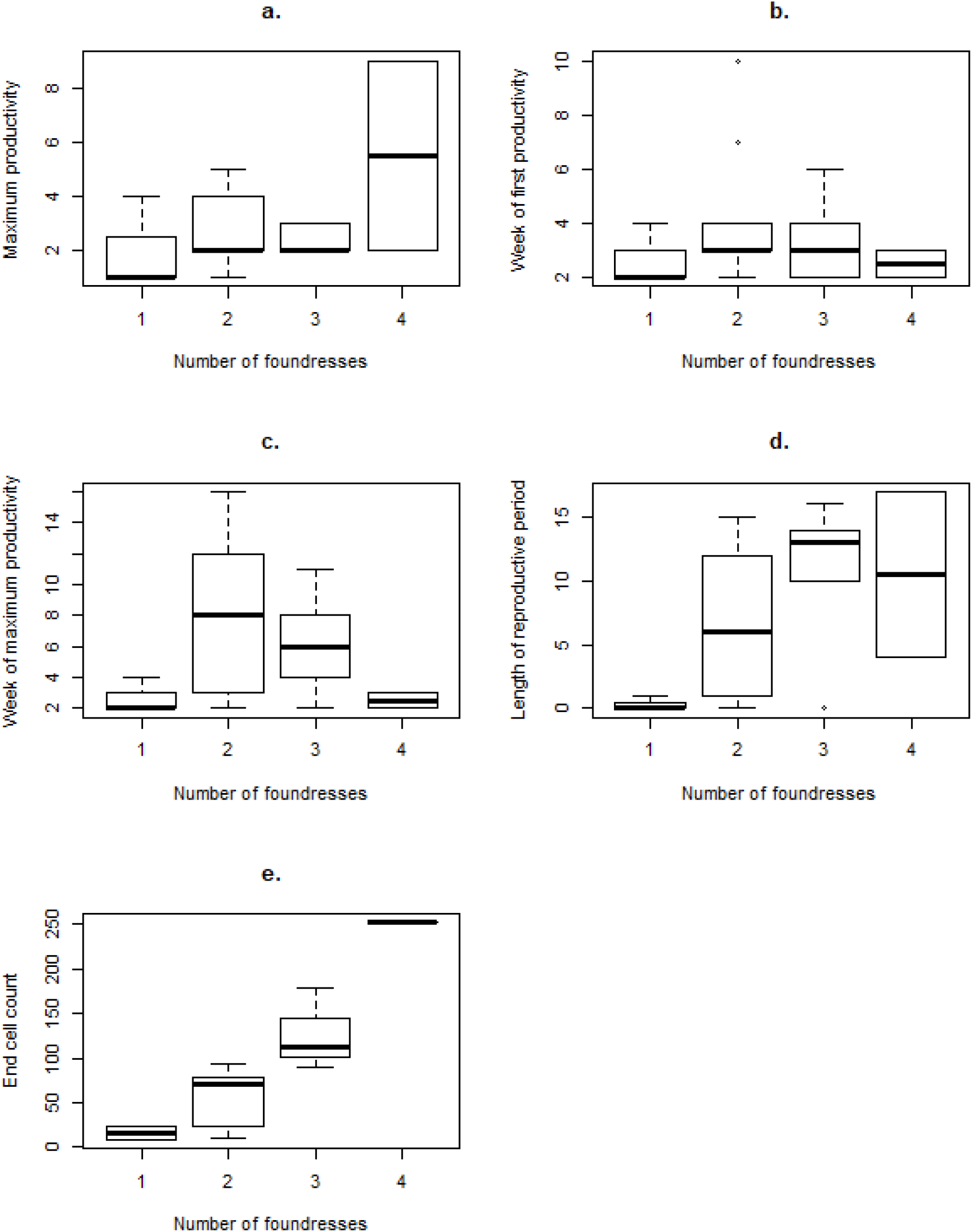
The relationship between the number foundresses in a *P. fuscatus* colony, and each of the 5 life-history traits. We have removed the single colony with five foundresses from this plot. a. Maximum productivity (i.e., the maximum number of new cells produced in any one week), b. Week of first productivity, c. Week of maximum productivity, d. Length of reproductive period, e. Final number of cells. Observations were started on June 12^th^.

In contrast to previous years (Wright, Tibbetts, McEwen, lichtenstein & Pruitt, in revision), parasitism by *Chalcoela iphitalis* moths was so infrequent as to prohibit its inclusion in our life-history syndrome analysis. Only 12% of nests harbored any parasitic moths at all. Nearly all of these colonies contained just one or two parasitic moths, although one *P. metricus* colony harbored an impressive 11 parasitic moths.

## DISCUSSION

Here we tested for the presence of life-history trade-offs and syndromes in paper wasps, whether they differed between species, and whether queen traits (size, boldness, coalition forming) might underlie between-colony differences in colony life-history. We detected no evidence of life-history trade-offs in either species of paper wasp considered here. Instead, we found that most colony performance metrics were positively correlated together for both species, but in subtly different species-specific manners. While we detected no evidence that queen boldness or body size contributed to differences in colony life-history in *P. metricus*, we found that cofoundress associations increased colony productivity in *P. fuscatus*, though not in a manner disproportional to foundress number.

Neither body size nor boldness appear to contribute significantly to colony life-history or performance in this population of *P. metricus*. Prior studies on *P. metricus* collected from this same field site found that bolder queens and larger queens were more productive than their small and shy counterparts (Wright et al., in revision). This raises the question of why, then, did we not detect such effects in this study. One possible explanation is that the prior study homogenized between-colony differences in microhabitat quality because it deployed nests within identical protective wooden boxes. This hints that the natural variation in nest site quality (i.e. temperature, light exposure, humidity, geometry, etc.) might drown out the signature of queen boldness or size. Studies in other species of social insect have recovered evidence for a colony-level pace-of-life (Reale, Garant, Humphries, Bergeron, Careau & Montiglio, 2010), where colonies can be placed along an axis of high-risk vs low-risk strategies that preferentially invest in reproductive output versus colony protection, respectively (Bengston, 2018; Bengston & Dornhaus, 2015; Bengston et al., 2017; Segev et al., 2017). More broadly, between-colony differences in collective aggressiveness and activity level have been associated with colony growth rates in several additional species (Blight et al., 2016; Bockoven et al., 2015). However, we recovered no evidence for such an association here. This is despite the fact that queen boldness is known to shape the collective aggressiveness of workers in *P. metricus* (Wright, Skinker, Izzo, Tibbetts & Pruitt, 2017). Thus, at least for now, we do have support for a pace-of-life syndrome in these species.

While we recovered no evidence of performance trade-offs between the life-history traits considered here, we detected strong evidence of life-histories syndromes that varied subtly between species. The model for a species-specific covariance structure between colony life-history traits had the highest support in our analysis. This is likely because the factors that underlie between-colony differences in colony life history vary considerably by species. For *P. metricus*, the factors underlying this syndrome remain elusive, but they are likely to include individual differences in condition or traits (e.g., habitat choice) not evaluated here. However, for *P. fuscatus*, between-colony differences in the syndrome are associated with variation in foundress number, which exhibits considerable intraspecific variation in our focal population.

In *P. fuscatus*, colonies with multiple queens exhibit higher performance across the entire constellation of life-history traits considered here, and in an apparently linear *per capita* manner. In other words, these differences vanish when we reanalyse colony life-history on a per foundress basis. Thus, while there appears to be a colony-level benefit to foundress coalitions, the payoff does not appear to be disproportionately large with the addition of the first helper, nor does the payoff appear to diminish with helpers 2-3. This is in contrast to the usual finding that the per capita benefit of adding cofoundresses decreases with increasing coalition size (Bono & Crespi, 2008; Jerome, McInnes & Adams, 1998; Karsai & Wenzel, 1998; Soucy, Giray & Roubik, 2003). Given these payoff structure, and the fact that foundress coalitions are not composed of clones, questions arise as to why joiners ever join such coalition (Bernasconi & Strassmann, 1999). We conjecture that either (1) such foundress coalitions might buffer colonies from total failure (e.g., via usurpation) in particularly stressful years or habitats (Tibbetts & Reeve, 2003), and that our current snapshot merely fails to capture harsh enough conditions, or (2) that only wasps of low condition/quality form coalitions, as it enables them to achieve a per-capita reproductive output that would not be possible if they founded a nest alone (Nonacs, 2001). Thus, coalition formation would represent a condition-dependent strategy. These hypotheses are not mutually exclusive but require longer-term or more in-depth data than we possess here to be fully excluded.

Intraspecific variation provides evolutionary biologists a spectrum with which to test for associations between traits and performance outcomes. Although there is a large and expanding literature devoted to documenting associations between individual variation behaviour and life-history traits (e.g., the pace-of-life literature) our data do not support a link between one of the most commonly evaluated behavioural traits (boldness) and life-history in paper wasps. Instead, intraspecific variation in foundress coalitions, which are known to have performance benefits for many kinds of social insect (ants: Bartz & Holldobler, 1982; thrips: Bono & Crespi, 2008; e.g., wasps: Tibbetts & Reeve, 2003), was the only predictor of between-colony differences in performance. Future work should target whether the tendency to join such coalitions is itself a diagnostic feature of an individual and, if so, whether it might be locked together with other kinds of individual differences reminiscent of a pace-of-life syndrome or some yet-identified conceptual framework.

## Acknowledgements

We are indebted to The University of Pittsburgh’s Pymatuning Laboratory of Ecology for permitting us to conduct these studies on their land, as well as the G. Murray McKinley Research Fund and the Arthur and Barbara Pape Endowment Award research grants provided through the University of Pittsburgh to CMW. Funding for this work was provided by the Tri-agency Institutional Programs Secretariat Canada 150 Chairs Program to JNP.

